# Forecasting Human Sleep Dynamics with a Latent-State Framework

**DOI:** 10.64898/2026.11.23.768504

**Authors:** Guokun Zhang, Luming Li, Huiling Yu

## Abstract

Sleep is essential for health, and forecasting its evolution could clarify its temporal organization and guide intervention timing. We developed a representation-forecasting model using electroencephalography from 5,419 participants. From 30-minute histories, 10-minute forecasts achieved 90.71% four-class accuracy in the 10% of windows with lowest predicted risk and averaged 72.11% overall. Direction-specific encoder feature axes had spectral correlates. Forecast changes under fixed-strength perturbations along these axes can quantify window-specific sensitivity varying with observed transition destination and bout timing. Physiological validation linked greater simulated perturbation sensitivity to stronger observed sleep-state responses following acoustic stimulation and respiratory events. Clinical profiling linked older age and higher apnoea-hypopnoea index to stronger modeled Wake-directed and weaker N3-directed responses. These results demonstrated that sleep-state forecasting could quantify dynamics, prioritize candidate intervention windows, and help interpret post-intervention changes.

## Introduction

Sleep is a fundamental physiological process of human beings and chronic sleep disturbances are associated with multiple health problems[1], [2]. As sleep is a dynamic process characterized by transitions among distinct brain states, understanding the internal temporal organization is central to sleep modulation[3], [4]. Forecasting the subsequent state transitions from recent sleep history therefore offers a quantitative approach to investigating these relationships and inform the selection of optimal intervention windows.

Theoretical models and empirical studies have provided evidence for the predictive temporal structure of sleep. The two-process model demonstrated the influences of homeostasis and circadian rhythms on sleep [5]. At the stage-sequence level, Yetton et al. used Bayesian networks to predict subsequent stages from preceding stages and durations [6]. Moving beyond discrete labels, sleep-onset trajectories in EEG feature space were quantified, revealing dynamics consistent with a fold bifurcation and critical slowing before the transition [7]. NREM-to-REM transitions were also be predicted up to 4 min ahead from EEG spectra, although predictions of the reverse transition did not outperform a time-in-stage baseline [8]. More recently, Lozano et al. showed that brainstem population activity in mice predicted both upcoming transition destinations and responses to REM-promoting stimulation [9]. Together, these studies reveal temporal dependencies in sleep architecture and physiological signals that anticipate specific state transitions. However, how well recent sleep history predicts future sleep states across forecast horizons and over the course of the night remains incompletely characterized. Addressing this gap could provide a quantitative basis for identifying candidate intervention windows.

Here, we developed a framework that jointly learns EEG representations and forecasts latent-state trajectories from recent sleep history. Using whole-night PSG data from 5,419 Sleep Heart Health Study (SHHS) participants [10], we quantified forecasting performance across sleep states, forecast horizons, times of night and evaluated cross-cohort generalization. Attribution and perturbation analyses identified direction-specific EEG features and characterized how simulation-derived perturbation sensitivity varied with transition context and timing. We further validated model outputs using data from acoustic stimulation and respiratory events. This framework characterizes the state and temporal dependence of sleep predictability and could inform the selection of candidate intervention windows.

## Results

### Sleep-state forecasting with time-dependent performance

We first developed a joint representation-forecasting framework comprising an EEG epoch encoder and a sequence predictor (Fig. 1A). The encoder mapped each 30-seconds EEG epoch to a 256-dimensional embedding, termed an EEG token, and projected it into a two-dimensional latent state, with the sequence predictor forecasting the subsequent latent-state trajectory from preceding EEG tokens. Representative windows illustrated the comparison between observed and forecast latent trajectories and sleep stage sequences over 10-min forecast horizon (Fig. 1B). For 10-minute-ahead predictions using 30 minutes of preceding sleep history, the model achieved a mean four-class sleep-stage prediction accuracy of 90.71% (95% CI, 90.08–91.32%), with a corresponding mean latent-state relative absolute error (RAE) of 0.261 (95% CI, 0.254–0.268) in the 10% of windows with the lowest predicted risk of sleep-stage error (Fig. 1C; see Supplementary Fig. S1C for predicted versus observed sleep-stage error rates). Across all forecast windows, mean sleep-stage accuracy and latent-state RAE were 72.11% (95% CI, 71.75–72.49%) and 0.404 (95% CI, 0.400–0.409), respectively (Fig. 1D), with the model outperforming the no-change baseline on both metrics (Supplementary Fig. S1A,B).

**Fig 1.**
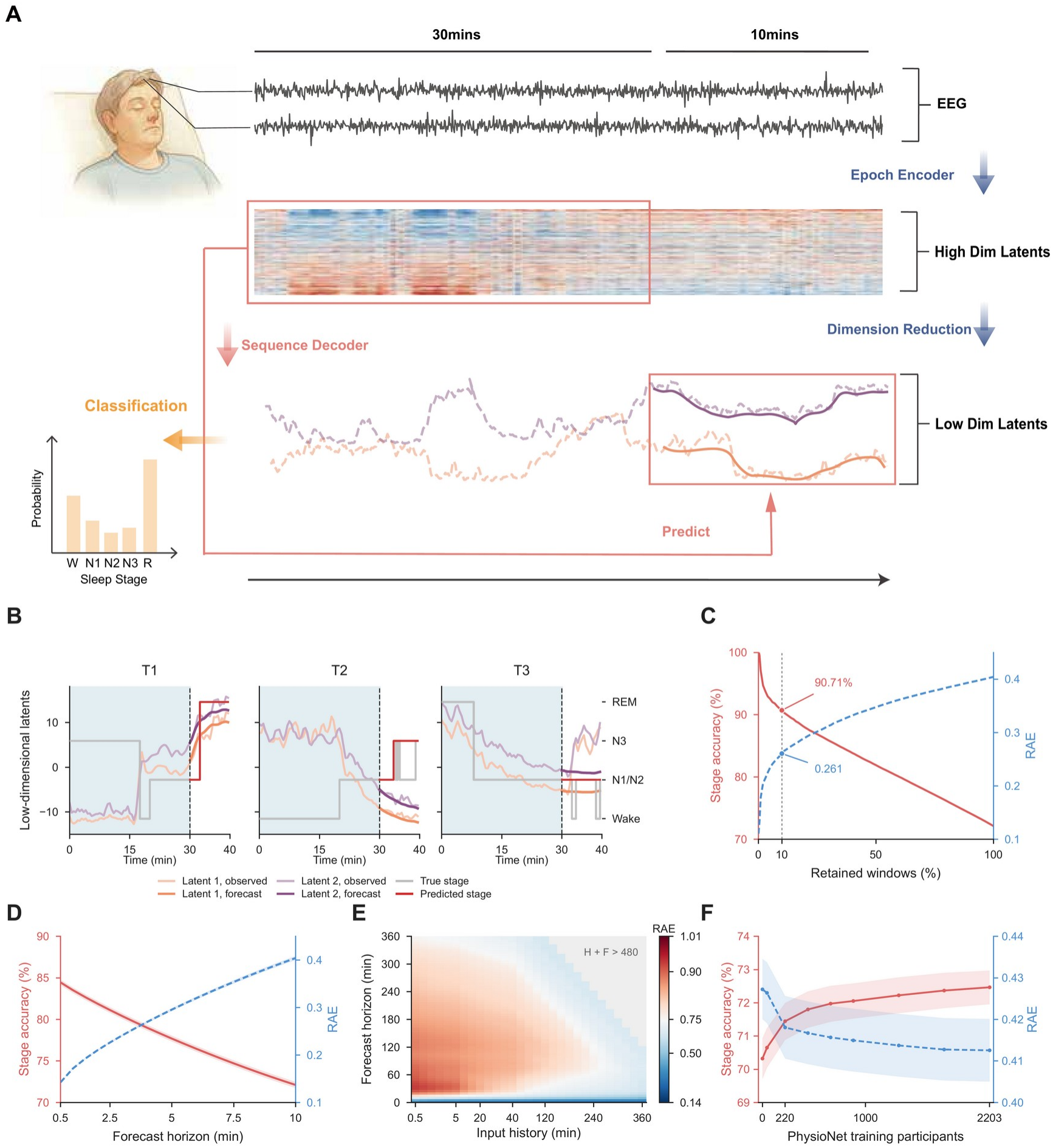
Sleep forecasting performance across windows and forecast horizons. (A)EEG epoch encoding and forecasting of sleep-state trajectories. N1 and N2 are pooled for four-class sleep staging in C, D and F. (B) Three example windows showing 30 min of EEG history (shading) and the subsequent 10 min of observed and forecast latent states and sleep stages. (C) Four-class accuracy (solid red, left axis) and latent-state relative absolute error (RAE, dashed blue, right axis) as increasing fractions of windows are retained. Windows were ranked by the mean predicted stage-error probability across the 20 forecast epochs. Metrics were averaged within each retained night and then across nights with equal weight. Dots mark the lowest-risk 10% of windows (91,664 windows from 1,058 nights). The 95% confidence intervals (CIs) reported for the retained subsets were obtained by resampling nights 2,000 times. (D) Cumulative accuracy and RAE from the forecast origin through each horizon. Lines show means across 1,084 SHHS test nights and shading shows pointwise 95% night-bootstrap CIs. Panels B–D use the fixed 30-min-input, 10-min-output model. (E) Endpoint RAE of the variable-length model for 1,921 tested combinations of 29 input-history lengths and 77 forecast horizons. Each combination uses its own eligible windows, averaged within nights and then equally across nights. Gray cells exceed 480 min of combined history and forecast. (F) External validation on the same 741,856 windows from 943 held-out PhysioNet participants in all nine conditions. Zero training participants denotes direct transfer of the SHHS-trained model. For the other conditions, the sequence predictor and stage readout were adapted while the encoder remained frozen. Lines show participant-equal means for accuracy (red) and RAE (blue). Shading shows pointwise 95% CIs from 10,000 paired participant-bootstrap resamples.

We next examined how input-history length and forecast horizon influenced endpoint forecasting performance. At a fixed forecast horizon, longer input-histories were generally associated with lower endpoint RAE. When input-history length was held constant, endpoint RAE initially increased and then decreased as the forecast horizon lengthened, with the turning point varying with input-history length (Fig. 1E). We then examined within-night variation under a fixed prediction setting, using a 30-min input history and a 10-min forecast horizon. Under this setting, endpoint RAE was lower in the second half of the recording than in the first (0.5363 versus 0.6428, Supplementary Fig. S1D). Together, these findings suggest that longer input histories provide useful predictive context, while endpoint forecasting performance varies with both forecast horizon and position within the night.

Having characterized forecasting performance across input-history lengths and prediction horizons, we assessed cross-cohort generalizability by appling the SHHS-trained model to 943 held-out participants from the PhysioNet 2026 Challenge [11], [12]. Without training on PhysioNet data, the model achieved a mean four-class accuracy of 70.32% and a mean latent-state RAE of 0.427 over the 10-min forecast using 30-min input-history length. Adapting the sequence predictor and stage readout while keeping the encoder frozen yielded additional improvements with increasing amounts of target-cohort data (Fig. 1F). These results support the model’s capacity for cross-cohort generalization and further adaptation.

### Direction-specific EEG features shape sleep-transition forecasts

We next examined whether sustained sleep-stage transitions could be forecast from preceding EEG. With a 30 minutes of input-history and a 10-min forecast horizon, the model distinguished upcoming transitions from continued stable sleep with a mean night-level AUC of 0.895 (95% CI, 0.884–0.906). Discrimination was stronger for the five sleep-to-sleep transiton types(AUCs, 0.905-0.987) than for the three sleep-to-Wake transition types (AUCs, 0.678-0.76, Supplementary Table S1).

To identify EEG features associated with forecasts toward specific target stages, we applied principal component analysis (PCA) to the model’s two-dimensional EEG representations. For each sleep stage, we defined a centroid as the mean position of its EEG representations in PCA space. We used the distance from each predicted representation to the target-stage centroid to quantify target-stage proximity, with smaller distances indicating forecasts closer to that stage (Supplementary Fig.S2 A-B). Integrated gradients (IG) [13] were used to identify individual 3-s input EEG frames with positive or negative contributions to predicted target-stage proximity, as illustrated for one N2⟶N3 window and four selected frames (Fig.2 A-B). For each of the eight transition directions, the transition-specific feature axes were constructed from normalized differences between CNN features extracted from positive- and negative-IG frames. In fivefold cross-validation grouped by night, these axes distinguished positive-from negative-IG frames, with mean night-level AUCs of 0.782–0.852 (all P ≤ 2.62 × 10^−8^ versus AUC = 0.5). For each transition direction, the corresponding axis yielded an AUC 0.125–0.553 higher than the mean AUC of axes for alternative destinations from the same source stage, supporting the direction specificity of these model-derived feature axes (all P < 0.001; Fig. 2C and Supplementary Table S2 A-C).To characterize the spectral correlates of these features, we compared EEG spectral power between the IG-defined frame groups. Compared with negative-IG frames, positive-IG frames showed higher relative delta power for N2⟶N3, lower delta and sigma power for N2⟶REM, and higher beta power for all three Wake-directed transitions (Fig. 2D and Supplementary Table S3A). Significant frequency clusters for each transition direction are listed in Supplementary Table S3B.

**Fig 2.**
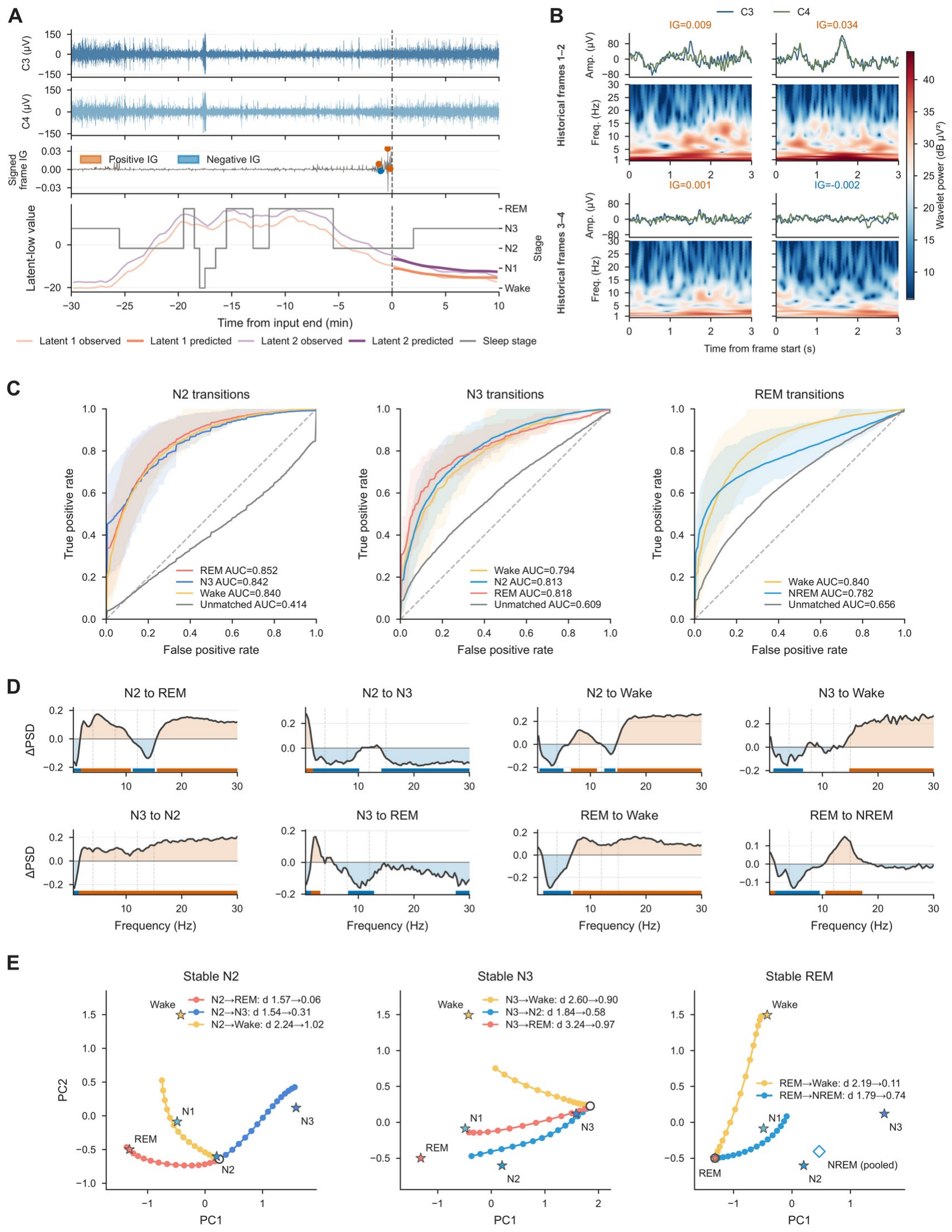
Direction-specific EEG features shape forecasts of sleep-stage transitions. (A) C3/C4 EEG, signed frame-level integrated gradients (IG), observed and predicted latent trajectories, and scored sleep stage in an illustrative N2⟶N3 window. The scored transition occurs 2 min after the forecast origin (time zero). The IG target is the negative mean standardized distance to the N3 centroid during the first 5 forecast minutes. (B) Four 3-s frames from A and their wavelet spectra. In reading order, the frames begin 76, 26, 12 and 66 s before the forecast origin. Numbers above the traces give signed IG values, and all four spectra share a power scale. (C) Fivefold cross-validation grouped by night tested whether CNN feature axes distinguish positive-from negative-IG frames. Colored curves use the axis matched to each destination. Gray curves summarize the alternative axes from the same source stage. Lines show mean nightly receiver operating characteristic curves, shading spans the nightly interquartile range, and labels report mean nightly areas under the curve (AUCs). Of 3,758 transition windows, 3,523 provided eligible frames of both signs. Two-sided Wilcoxon signed-rank tests compared nightly matched-axis AUCs with 0.5 and with the corresponding mean alternative-axis AUCs. Holm correction was applied across eight directions separately for each comparison family. (D) Night-averaged differences in log_10_ relative EEG power between positive- and negative-IG frames. Orange and blue indicate higher and lower power in positive-IG frames, respectively. Bars mark frequency clusters significant at P < 0.05 in a two-sided maximum-cluster-mass test with 10,000 night-level sign flips and a cluster-forming threshold of P < 0.05. Family-wise error was controlled across frequencies within each direction. (E) Mean forecast position during the first 5 forecast minutes after perturbing CNN features along each axis in the last 5 input minutes (α = 0–15). The observed stage remained N2, N3 or REM for 5 min on either side of the forecast origin. Dots mark perturbation strengths, open circles mark α = 0, stars mark stage centroids, and the diamond marks the pooled NREM centroid. Labels report distances from the mean forecast position to its target at the lowest and highest strengths.

We next tested whether perturbations along these CNN feature axes and corresponding spectral perturbations of raw EEG could shift model forecasts toward the target stages. We selected N2, N3 and REM windows in which the observed stage remained unchanged for 5 min before and 5 min after the forecast origin. We perturbed CNN features from the final 5 min of input along the corresponding feature axis (α = 0–15, Fig. 2E) and separately applied templates derived from the spectral differences between positive- and negative-IG frames to the final 5 min of raw EEG while preserving Fourier phase. In the model forecasts, both approaches generally reduced target-centroid distance as perturbation magnitude increased, with increases in target-stage proportion for CNN feature perturbations and target-region occupancy for raw-EEG perturbations. However, N3⟶N2 showed nonmonotonic responses under both approaches, with target-directed effects weakening at higher magnitudes (Supplementary Fig. S3). Together, these findings identified direction-specific, model-derived EEG features with interpretable spectral correlates and showed that targeted perturbations could steer predicted sleep-state trajectories toward the corresponding stages.

### Perturbation gains reflect observed sleep responses and vary with transition destination and timing

Having shown that direction-specific perturbations could alter predicted sleep-state trajectories, we next examined which windows yielded larger target-directed gains under a fixed perturbation direction and strength, and how these gains varied with subsequent transition destination and progress through source-stage bouts. For each eligible N2, N3 or REM window throughout the night, we applied each source-specific perturbation separately to the CNN features of the final 30-s input epoch at a fixed strength. We compared the perturbed and unperturbed forecasts over the same subsequent 10 min. Distance gain (ΔD) measured the reduction in mean distance to the target-stage centroid in PCA space. Occupancy gain (ΔP) measured the increase in the fraction of forecast epochs falling within the target-stage region. Larger gains indicated greater sensitivity of the model forecast to the applied perturbation. A whole-night example showed substantial variation among windows within the same sleep stage (Fig. 3A).

**Fig 3.**
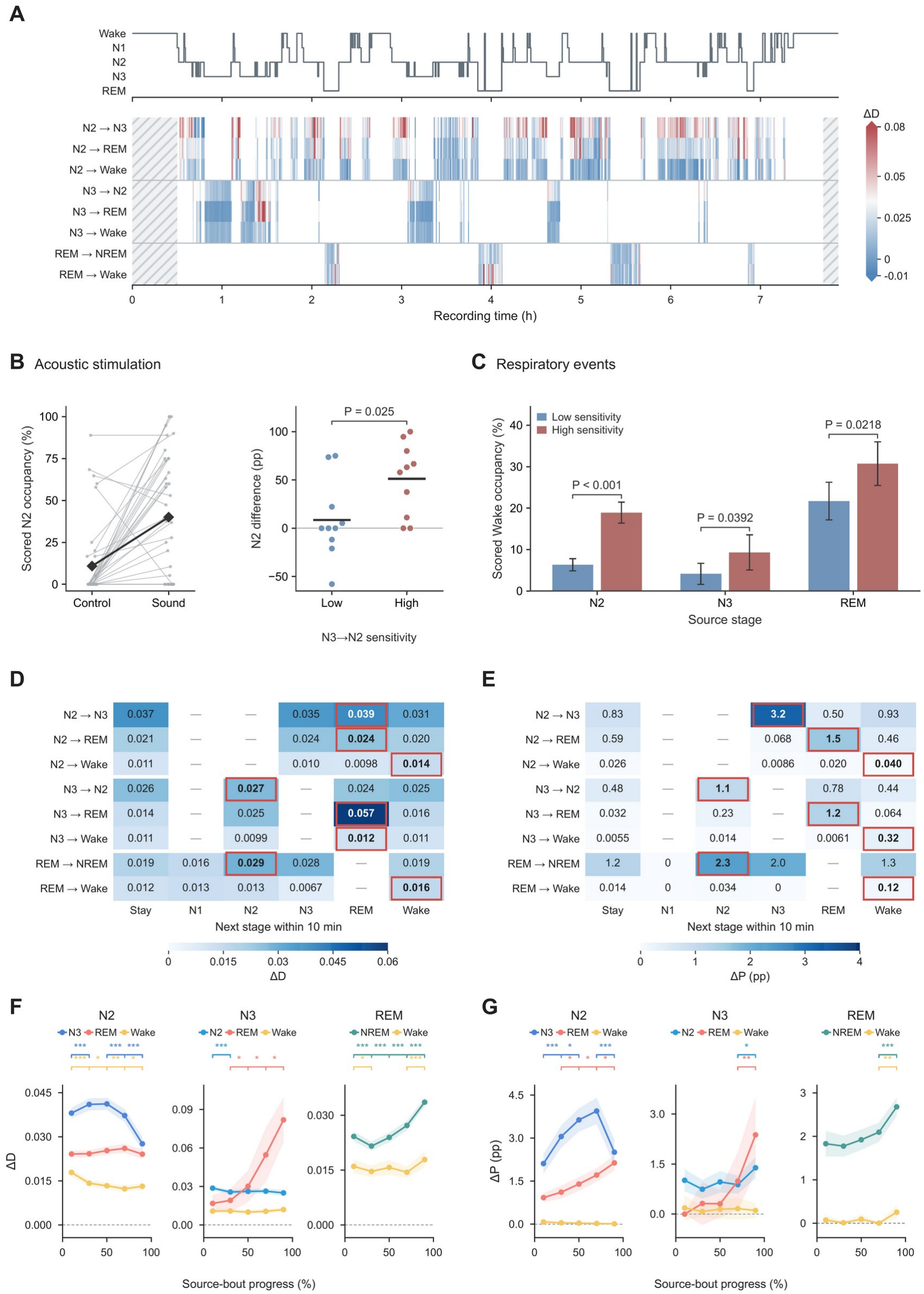
Direction-specific perturbation gains reflect observed sleep responses and vary with transition destination and timing. (A) Scored stages and distance gains in one night. CNN features from the final 30-s input epoch were perturbed at α = 1. Distance gain (ΔD) is the reduction in mean target-centroid distance over the next 10 min relative to the unperturbed forecast. Occupancy gain (ΔP) is the increase in target-region occupancy in percentage points (pp). Hatching indicates insufficient data. (B) Manually scored N2 occupancy after acoustic stimulation in N3. Lines connect each stimulation window to its control from the same participant’s unstimulated night (39 pairs, seven participants). Diamonds show means. Follow-up used complete aligned epochs after the onset epoch until 10 min or the next sound train. At right, points show stimulation-minus-control differences for the lowest and highest quartiles of prestimulation N3⟶N2 ΔD (10 pairs each). Lines show means. Groups were compared by a two-sided Welch t test. (C) Manually scored Wake occupancy in the 19 complete epochs (9.5 min) after the respiratory-event onset epoch. Of 3,907 hypopnea or apnea events, 1,956 fell in the top or bottom quartiles of Wake-directed ΔD within stage (657 per group in N2, 117 in N3 and 204 in REM). Bars show event means and pointwise 95% t CIs. Two-sided Welch t tests compared groups within each stage, with Holm adjustment across stages. (D, E) Mean ΔD (D) and ΔP (E) across windows for each perturbation (rows) and subsequent stage (columns). Stay denotes an unchanged stage over the next 10 min. Transition groups required a direct transition within those 10 min between bouts lasting at least 5 min each. Red outlines mark row maxima before rounding. Dashes indicate empty groups. (F, G) ΔD (F) and ΔP (G) across five equal intervals from the first to the last 30-s epoch of the source-stage bout (3,727 transitions, 1,028 nights). Curves show equally weighted nightly means and pointwise 95% t CIs. Brackets mark two-sided paired t tests on nightly means in adjacent intervals, with Benjamini–Hochberg correction across the four comparisons within each curve. One, two and three stars denote adjusted P < 0.05, 0.01 and 0.001, respectively.

We first tested whether simulation-derived ΔD was associated with window-specific sleep-state responses following external acoustic stimulation and endogenous respiratory events. In an independent public acoustic-stimulation dataset [14], we analyzed 39 N3-stage stimulation windows from seven participants, each paired with a matched control window from the same participant’s unstimulated night.The stimulation protocol was intended to suppress slow-wave sleep and promote shallower sleep. Mean manually scored N2 occupancy was 40.05% in post-stimulation windows and 10.86% in control windows. We then stratified the 39 stimulation windows by prestimulation N3⟶N2 simulated ΔD, designating the upper and lower quartiles as high- and low-sensitivity groups, respectively. We quantified the subsequent stimulation responses as the paired stimulation-minus-control difference in N2 occupancy. This difference was greater in the high-sensitivity group than in the low-sensitivity group (51.11% versus 8.55%, P = 0.0253, Fig. 3B). We next examined whether Wake-directed simulated ΔD was associated with observed Wake occupancy following naturally occurring respiratory events in SHHS dataset. We included 3,907 eligible hypopnea and apnea events. Within each source stage, windows in the upper and lower quartiles of Wake-directed simulated ΔD were classified as high- and low-sensitivity windows, respectively. Manually scored post-event Wake occupancy was higher in the high-sensitivity group than in the low-sensitivity group across the N2, N3 and REM source stages: 18.91% versus 6.33%, 9.31% versus 4.14%, and 30.73% versus 21.70%, respectively (Holm-adjusted P = 2.48 × 10^−16^, 0.0392 and 0.0218 across the three comparisons, Fig. 3C). Together, these findings support simulated ΔD as a model-derived indicator of window-specific perturbation sensitivity to sleep-state modulation.

Under a fixed perturbation direction and strength, perturbation sensitivity also varied with the window’s subsequent observed stage trajectory. We classified windows as Stay if the scored stage remained unchanged throughout the next 10 min, and assigned windows to transition groups if a direct transition occurred within that period and the source and destination stages each persisted for at least 5 min. Across all eight perturbation directions, mean ΔP was highest in windows whose observed destination matched the perturbation target (Fig. 3E). For instance, N2⟶N3 perturbations produced a mean gain of 3.22% in windows that subsequently entered N3 while 0.83% in Stay windows. For N2⟶N3 and N3⟶Wake perturbations, however, ΔD was greatest in windows that subsequently entered REM (Fig. 3D).

Moreover, the relative timing of peak perturbation sensitivity differed across transition directions. We normalized progress through each source-stage bout preceding a sustained transition from 0 to 1, divided the bout into five equal intervals and averaged gains first within nights and then across nights with equal weight (Fig. 3F,G). Based on ΔP, N2⟶N3 was most sensitive during the 60–80% interval, when mean gain peaked at 3.94% before declining to 2.51% in the final interval (a decrease of 1.44%; P = 2.51 × 10^−6^; Fig. 3G). For N2⟶REM, N3⟶REM and REM⟶NREM, ΔP increased later in the bout, with the highest mean gains occurring in the final 80–100% interval. N3⟶N2 and REM⟶Wake also showed their highest mean ΔP in the final interval. Mean ΔD was highest in the 80–100% interval for N3⟶REM, N3⟶Wake, REM⟶NREM and REM⟶Wake; for example, N3⟶REM ΔD increased from 0.0168 in the first interval to 0.0819 in the last (Fig. 3F). For N2⟶N3 and N2⟶REM, mean ΔD was highest in the 40–60% and 60–80% intervals, respectively, whereas the highest means for N2⟶ Wake and N3⟶ N2 occurred in the 0–20% interval. These patterns suggested that potential intervention windows differed across transition directions and that the timing of greatest sensitivity also depended on the outcome metric.

### Forecast error and perturbation sensitivity vary with age and sleep-disordered breathing severity

Having characterized forecast error and linked perturbation sensitivity to observed responses to acoustic stimulation and respiratory events, we next used age and apnoea-hypopnea index (AHI) as illustrative characteristics to demonstrate the clinical variation in forecast error and model-estimated perturbation sensitivity in SHHS.

In a model including age, sex, body mass index (BMI), AHI and overnight latent-state mean absolute deviation (MAD),older age was associated with higher whole-night RAE (standardized β = 0.129, P = 1.24 × 10^−4^; Fig. 4A),whereas AHI was not significantly associated with whole-night error (β = −0.032, P = 0.452; Fig. 4B). Transition-specific analysis revealed different patterns: older age was associated with higher error at N2⟶REM (standardized β = 0.115, P = 0.0418), while higher AHI was associated with lower error at REM⟶Wake (β = −0.227, P = 0.00272; Fig. 4C).

**Fig 4.**
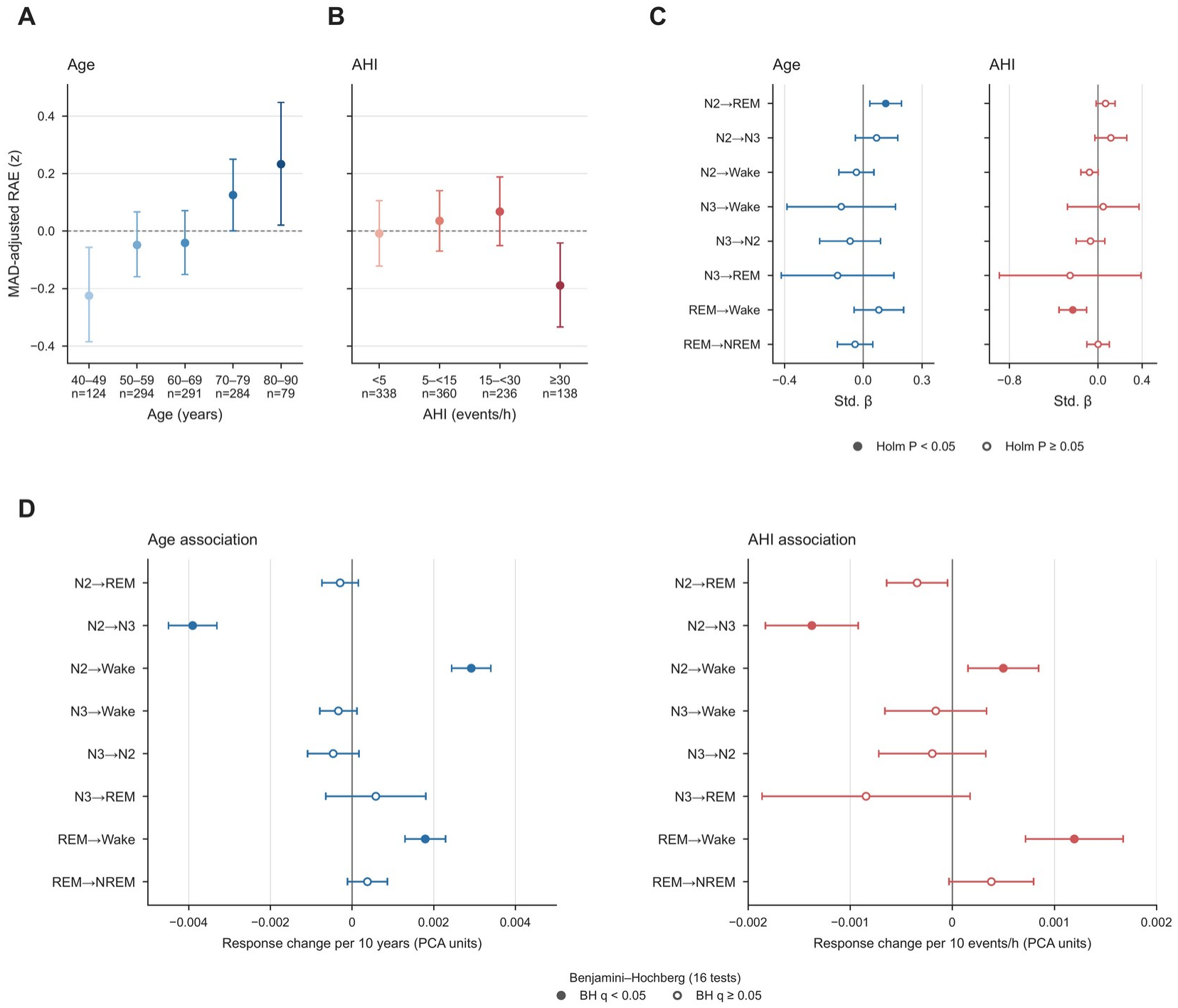
Associations of forecast error and directional perturbation sensitivity with age and sleep-disordered breathing severity. (A, B) Mean whole-night relative absolute error (RAE) by age (A) and apnoea–hypopnoea index (AHI, B) in 1,072 SHHS test nights with complete clinical data. RAE was adjusted for the overnight mean absolute deviation of each latent coordinate using coefficients estimated on 643 validation nights, then standardized in the test cohort. Error bars are 95% percentile CIs from 10,000 within-group night-bootstrap samples. Continuous age and AHI effects reported in the text were estimated in an ordinary least-squares model including age, sex, body mass index and AHI, with HC3 standard errors and Holm adjustment across four covariates. (C) Standardized coefficients for age and AHI in regressions of transition-specific RAE on the same four covariates. Errors from 3,725 eligible transitions were averaged within night and direction before fitting. Panels A–C use unsmoothed latent targets. Horizontal bars show pointwise 95% HC3 Wald CIs. Filled markers indicate P < 0.05 after Holm correction across the eight directions for each exposure. (D) Change in distance gain (ΔD) per 10 years of age or per 10 AHI events/h. Features of the final 30-s input epoch were perturbed at α = 1 in windows with the same source stage for 5 min on either side of the forecast origin. Gains were averaged over the 10-min forecast and then within each night and direction. Separate unadjusted regressions were fitted for age and AHI (737–1,065 nights per direction). Bars show pointwise 95% t CIs using HC3 standard errors. Filled markers indicate Benjamini– Hochberg-adjusted P < 0.05 across all 16 exposure-by-direction tests. The separate severity-group contrasts reported in the text compared AHI ≥30 with <5 events/h in unadjusted categorical models, with HC3 t inference and Benjamini–Hochberg correction across eight directions.

For model-estimated perturbation sensitivity, age and AHI showed concordant directional perturbation sensitivity. We applied feature perturbations of magnitude α = 1 along the eight established directions to the final 30 seconds of the input, with the simulated response quantified by ΔD. In stable N2 windows, both older age and higher AHI were associated with weaker N2⟶N3 and stronger N2⟶Wake responses. In stable REM windows, both were also associated with stronger REM⟶Wake responses (Fig. 4D). For severity-group comparisons, we used AHI categories of <5, 5–<15, 15–<30 and ≥30 events/h, with <5 events/h as the reference [15].

Consistent with the AHI-related associations described above, participants with AHI ≥30 events/h had weaker N2⟶N3 responses than those with AHI <5 events/h (difference, −0.00570; 95% CI, −0.00820 to −0.00319; multiplicity-adjusted P = 7.39 × 10^−5^). They also had stronger N2⟶Wake responses (difference, 0.00234; 95% CI, 0.00049–0.00418; adjusted P = 0.0346) and REM⟶ Wake responses (difference, 0.00520; 95% CI, 0.00269-0.00772; adjusted P = 2.13 × 10^−4^). These findings reveal clinical variation in forecast accuracy and perturbation sensitivity, with older age and higher AHI associated with greater model-estimated vulnerability to Wake-directed disruption.

## Discussion

The present findings support a conditional view of sleep predictability. Recent EEG history contains information about subsequent sleep-state evolution, but forecasting performance varies across temporal settings, transition directions and individuals. Direction-specific perturbations provide a complementary perspective by characterizing how predicted trajectories respond to targeted changes in the input. Together, these approaches offer a framework for examining both the predictability of spontaneous sleep dynamics and the potential to identify windows responsive to modulation.

### Conditional predictability of sleep dynamics

Sleep is partially predictable, with physiological mechanism imposing temporal constraints on its evolution. Homeostatic modulation links sleep intensity to prior sleep-wake history, while circadian rhythms shapes the timing of sleep and wakefulness [16], [17]. Previous studies have linked continuous EEG measures to imminent arousal, predicted NREM-to-REM transitions and future sleep-stage sequences [8], [18], [19]. These physiological and forecasting studies provide a basis for anticipating sleep evolution from its preceding history.

Several factors may contribute to the limited predictability of sleep. First, scalp EEG provides limited information about regional neural activity and autonomic function, so similar EEG histories may correspond to different underlying physiological states and subsequent sleep trajectories [20], [21], [22]. Secondly, Dynamical instability can amplify small differences in initial states, causing future trajectories to diverge. Human intracranial recordings have revealed state-dependent chaotic-like activity [23]. Additionally, stochastic models of arousal and sleep onset show how random fluctuations can trigger transitions and alter their timing [24], [25]. Thus, sleep dynamics is a predictable temporal structure while leaving individual transitions uncertain. In this study, we use this partial predictability to investigate sleep-state dynamics and systematically quantify how recent EEG history constrained subsequent evolution and how direction-specific input perturbations altered predicted trajectories. This extends previous work from predicting sleep events and stage sequences to generating testable hypotheses about sleep-state transition dynamics.

### Forecasting error and perturbation sensitivity as probes of sleep-state dynamics

Previous studies have established physiological descriptions of specific sleep transitions. Using simultaneous scalp and intracranial EEG, Peter-Derex et al. showed that NREM-to-REM transitions unfold asynchronously across cortical regions, with REM activity in the lateral occipital cortex emerging approximately 2 min before scalp-defined onset [26]. Li et al. represented wake-to-sleep progression as a trajectory in EEG feature space and identified fold-bifurcation dynamics with critical slowing before the transition tipping point, enabling real-time prediction of sleep progression [7]. Our study extends these analyses of particular transitions to eight transition directions from N2, N3 and REM within a shared representation learned directly from raw EEG. Destination-specific feature axes and their spectral correlates connect the model’s predictive features to measurable electrophysiological patterns. For example, features favouring N2-to-N3 forecasts were associated with higher relative delta power, whereas those favouring N2-to-REM forecasts were associated with lower delta and sigma power. This integrates physiological interpretation with trajectory forecasting across multiple destinations, characterizing how distinct EEG patterns within the same scored stage relate to different subsequent courses.

Perturbation analyses quantify the functional consequences of these feature changes within the forecasting model. Previous whole-brain models used simulated stimulation to investigate sleep–wake transitions [27], while mouse experiments demonstrated that ongoing brainstem activity gates responses to REM-promoting inputs [9]. Our study connects measurable EEG changes to their effects on trajectories conditioned on recent sleep history. Feature-space and raw-EEG spectral perturbations produced concordant target-directed forecast changes. The nonmonotonic N3-to-N2 response further showed that increasing a feature change along a target-related axis did not consistently strengthen the corresponding trajectory shift. These analyses establish quantitative relationships between the direction and magnitude of representation changes and their predicted consequences. Further studies should map these feature axes and perturbation magnitudes to experimentally measured changes in neural activity and EEG. Stimulation experiments should then determine which modalities and parameters induce the corresponding feature changes and whether the resulting sleep-state responses agree with the forecasts.

### Selection intervention windows and evaluating sleep modulation

Intervention timing is a determinant of how effectively a stimulus engages sleep physiology. Auditory stimulation aligned with slow-oscillation up states enhances sleep rhythms and memory, whereas out-of-phase stimulation is ineffective [4]. Similarly, closed-loop auditory stimulation enhances slow oscillations and spindle occurrence within specific phase windows, with a narrower effective window in older adults [28]. These findings highlight intervention timing as a key determinant of sleep modulation efficacy and indicate that conventional sleep staging alone provides limited guidance for selecting suitable intervention windows. Previous studies have investigated how ongoing physiological activity can inform stimulation timing. Prestimulus EEG can identify transient periods of behavioural responsiveness during sleep [29], and combined EEG and cardiac phase information has been associated with stronger auditory-evoked slow-wave responses [30]. However, these studies demonstrate physiological markers of responsiveness but do not systematically compare intervention windows across the night for different sleep-state transition targets. Our framework maps direction-specific perturbation sensitivity across the night, allowing candidate windows to be compared for a specified transition. These estimates could further guide window selection according to the intended sleep-state change.

Forecasting also provide a reference for evaluating whether modulation produces additional change. Recent optogenetic studies have evaluated sleep modulation through changes in transition probabilities, state occupancy and sleep-episode duration [9], [31]. A sham-controlled rTMS study in patients with insomnia assessed modulation effects using sleep onset latency, sleep efficiency and N3 duration [32]. However, these summary measures alone do not provide a window-specific reference trajectory for how an individual’s sleep would have evolved without intervention. Predicted untreated trajectories have been used as references in time-series intervention evaluation [33], [34], [35], including applications to biomedical sensor data and sleep-apnea monitoring [36], [37]. In our framework, unperturbed forecasts could provide a window-specific reference for assessing whether stimulation advances target-stage entry, increases occupancy or prolongs maintenance beyond expected spontaneous sleep evolution.

## Acknowledgments

We thank Prof. Yang Yuan from the Institute for Interdisciplinary Information Sciences, Tsinghua University, Prof.Jingying Ye and Prof. Guoping Yin from the Department of Otolaryngology-Head and Neck Surgery, Beijing Tsinghua Changgung Hospital, School of Clinical Medicine, Tsinghua University, for their helpful suggestions throughout this study.

## Materials and Methods

### Study design and data source

We used whole-night polysomnography data from Visit 1 of the Sleep Heart Health Study (SHHS) [10], including one recording per participant. Of the 5,793 recordings with matched EEG and sleep-stage data, 5,419 met the requirement of more than 2 h of largely artifact-free usable signal in each EEG channel (SHHS quality scores of at least 2). Participants were divided into training, validation, and independent test sets containing 3,685, 650, and 1,084 participants, respectively. All EEG epochs and forecast windows from a participant belonged to the same subset. We evaluated forecasting performance in an external cohort using the PhysioNet 2026 Challenge dataset [11], [12]. Of 6,620 available records, 6,293 participants with one recording each met the EEG, reference and human-staging requirements, including at least one continuously valid 40-min interval. Participants were split into training (n = 4,406), validation (n = 944) and test (n = 943) sets.

To test whether simulated perturbation gains corresponded to responses to acoustic stimulation, we used the public slow-wave sleep suppression dataset of Ukraintseva and Liaukovich [14]. The 19-channel EEG recordings included sound-onset markers. We analyzed 39 N3 stimulation windows from seven participants, each paired with a control window from the same participant’s unstimulated night.

### EEG and sleep-stage preprocessing

For SHHS and PhysioNet, signals were converted to μV and divided into contiguous, non-overlapping 30-s EEG epochs aligned with manual sleep staging. Each EEG epoch was resampled to 100 Hz using MNE’s FFT-based resampling [38]. Five labels were retained: Wake, N1, N2, N3, and REM. The SHHS model inputs were the C4–A1 and C3–A2 EEG derivations from the EDF files, in that order. For SHHS, the original N3 and N4 stages were merged as N3.

For PhysioNet recordings, the two inputs were matched to C4–M1 and C3– M2, in that order, using native bipolar derivations or re-referencing separate central and mastoid signals when their common reference was documented.

For the acoustic-stimulation recordings, C4–A1 and C3–A2 were reconstructed from the documented EDF channels and converted to μV using the EDF calibration. Each complete 30-s epoch was independently resampled from 250 to 100 Hz. Expert stage-onset and end annotations were aligned to the 30-s grid, assigning a stage only when it covered more than half of an epoch. Ambiguous or unpaired annotations were left unlabelled, and incomplete terminal epochs were discarded. The recorded amplitude scale was retained.

### Overall modeling framework

To forecast future sleep states from historical EEG, model training comprised two stages. First, the EEG epoch encoder and short-horizon forecasting module were trained jointly under sleep-stage supervision and future-state forecasting objectives. Each 30-s EEG epoch was represented as a 256-dimensional EEG token and a corresponding two-dimensional latent state. Second, the EEG epoch encoder was frozen, and whole-night recordings were converted in advance to EEG token sequences. An independent Transformer sequence predictor was then trained to forecast the next 10 min of two-dimensional latent states from the preceding 30 min of EEG tokens.

### EEG epoch encoder and joint representation learning

The EEG epoch encoder combined local frame features with temporal information across frames to construct a latent representation of each epoch. For EEG epoch *t*, the 30-s input from the two central channels was denoted *x*_*t*_ *∈* ℝ^2*×*3000^. Each EEG epoch was divided into overlapping 3-s EEG frames with onsets spaced 1 s apart, yielding 28 EEG frames. EEG frame *f*, denoted *x*_*t, f*_ *∈* ℝ^2*×*300^, was first processed by a multiscale one-dimensional residual convolutional network with kernel sizes of 3, 5, and 7 samples. The two channels were fused at this stage. Frame features were linearly mapped to 256 dimensions, supplemented with fixed positional encodings, and integrated within each EEG epoch by a four-layer, eight-head Transformer [39]. Attention pooling then yielded a 256-dimensional EEG token:

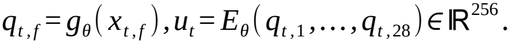

The EEG token was mapped to a two-dimensional latent state through a 256⟶128⟶2 projection head. The first linear layer was followed by batch normalization and an exponential linear unit (ELU) activation; the final layer was linear:

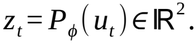

The two-dimensional latent state was also passed to a five-class linear head to obtain predicted probabilities for the five sleep stages:

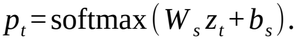

Joint representation learning used contiguous 30-min windows. The first 25 min (50 EEG epochs) provided the observed input; the final 5 min (10 EEG epochs) were replaced with learnable mask tokens in the short-horizon forecasting module. A two-layer, four-head Transformer short-horizon forecasting module used the observed EEG tokens to generate the ten future two-dimensional latent states in parallel. Sleep-stage targets used transition-aware soft labels. At the last EEG epoch before each stage transition, the one-hot target for the current stage was mixed with that of the immediately following stage with weights of 0.7 and 0.3, respectively. Hard labels were used at all other positions. The staging objective was soft-label cross-entropy across all 60 EEG epochs. The forecasting objective was the mean of the dimension-specific root-mean-square errors over the ten future positions, each normalized by the sample standard deviation of the corresponding true latent coordinate across the complete 60-epoch sequences in the same minibatch:

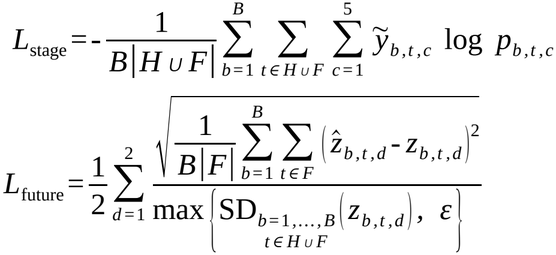

where *H* ={1,*…*, 50 } and *F* ={51,*…*, 60 } denote the historical and future epoch indices, respectively, and *B* is the local minibatch size on each GPU. For sample *b* at epoch *t*, 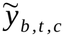 is the transition-aware target probability for sleep-stage class *c*, and *p*_*b,t,c*_ is the corresponding classifier output probability. The quantities *z*_*b, t, d*_ and 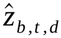denote the true and predicted latent coordinates in dimension *d*. The operator SD denotes the sample standard deviation across the indicated batch and time indices, and *ε* = 10^-6^ sets a lower bound on the normalization scale. The combined loss used for monitoring and validation was *L*_joint_ = *L*_stage_ + *L*_future_. We used PCGrad [40] to project conflicting task gradients before summing them with equal weights.

Training-window onsets were spaced 3 min apart. Both training stages used AdamW [41] with a fixed learning rate of 5 × 10^−5^, zero weight decay, bfloat16 mixed precision and gradient-norm clipping at 1.0, for a maximum of 50 training epochs. Joint representation learning used a global batch size of 160. Training stopped if the validation joint loss failed to improve for two consecutive training epochs. The checkpoint from the tenth training epoch, which had the lowest validation objective, was retained for subsequent analyses.

We selected a two-dimensional latent state because, across the same 650 validation nights, five-stage accuracy for the current epoch improved from one to two dimensions and showed no further improvement with a third. The third principal component accounted for less than 0.3% of total variance in the three-dimensional encoder (Supplementary Fig. S4A–D).

### Sleep latent-state sequence predictor

The sequence predictor generated future two-dimensional latent-state trajectories from historical EEG features. It used contiguous 40-min samples. The first 30 min contained 60 observed 256-dimensional EEG tokens, and the final 10 min contained 20 positions to be forecast. All future positions were replaced with learnable mask tokens. The model generated 20 future two-dimensional latent states simultaneously in one forward pass, implementing parallel, non-autoregressive forecasting. True future EEG, sleep stages, and latent states were not provided as model inputs. Denoting the sequence predictor by *F*_*ω*_, we obtained

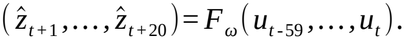

The sequence predictor had an internal dimension of 256 and comprised a four-layer, four-head Transformer. A linear output layer mapped its representations to two-dimensional latent states. The training objective was mean absolute error between the forecast states and the corresponding two-dimensional target states. Targets were smoothed with a centered moving average over three EEG epochs within each contiguous valid segment of each recording. Historical EEG tokens and forecast outputs were not smoothed. No auxiliary high-dimensional reconstruction loss was used. For a minibatch of *B* windows, the objective was

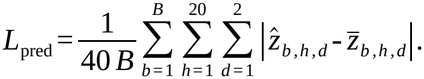

where 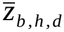 denotes the smoothed target and 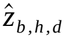 denotes the forecast value. The loss was averaged over windows, future positions and the two latent dimensions.

At this stage, the EEG epoch encoder was frozen. Training used all windows meeting the duration requirements, enumerated at 30-s intervals. During validation and testing, all eligible windows were enumerated at the same 30-s stride. The sequence predictor used a batch size of 800 and early stopping.

### Forecast error of two-dimensional latent-state trajectories

Model performance was evaluated using relative absolute error (RAE) in the two-dimensional latent state. For latent-state dimension *d*, a common scale was defined by the mean absolute deviation of all true future targets in the independenttest set. Let *z*_*i,d*_ and 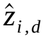denote the true and forecast values, respectively, and let *μ*_*d*_ denote the mean of the true future targets for that dimension:

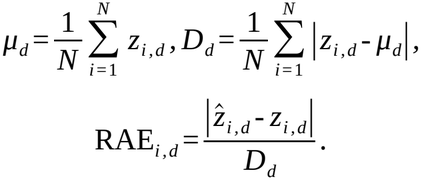

To reduce local random fluctuations in the latent-state targets, true trajectories were smoothed using a centered moving average over three EEG epochs.

### Sleep-stage decoding from forecast latent states

To test whether forecast trajectories retained sleep-stage information, N1 and N2 were merged a priori because their representations were similar. The resulting four classes were Wake, N1/N2, N3, and REM. The decoder received the complete 20-point forecast trajectory of two-dimensional latent states. It comprised a two-layer bidirectional long short-term memory network (BiLSTM) with a hidden dimension of 128 in each direction and inter-layer dropout of 0.15. Layer normalization, dropout, and a four-class linear layer followed the BiLSTM.

The decoder was trained exclusively on forecast windows from the validation set. The number of training epochs was selected using an internal 80%/20% participant-level split of the 650 validation participants, with 520 participants for fitting and 130 for tuning. The epoch count was selected by pooled four-class tuning accuracy; the decoder was then refitted on all 650 participants. The independent test set was not used for parameter selection. Each class was evaluated using one-versus-rest receiver operating characteristic (ROC) analysis. Macro-averaged area under the ROC curve (AUC) included only nights for which AUC was defined for all four classes.

### External validation

To assess cross-dataset generalization, we compared direct transfer of the SHHS model to the external test set with target-domain fine-tuning. External validation used a separate SHHS source-training run with 4,606 training and 813 validation participants. The sequence predictor and four-class readout were then refitted on all 5,419 SHHS participants for the numbers of training epochs selected within SHHS. The encoder remained frozen during fine-tuning, which used learning rates of 10^−5^ for the predictor and 10^−4^ for the readout.

### Baseline models

To assess forecast gains over persistence of the current state, the no-change baseline extended the last unsmoothed observed two-dimensional latent state throughout the forecast interval. Its sleep-stage baseline decoded this state with a multilayer perceptron (MLP), fitted exclusively to true two-dimensional latent states in the validation set. The resulting hard label was then extended throughout the forecast interval.

To compare the model with forecasts based on conventional spectral features, we included a power spectral density (PSD) baseline. For this baseline, absolute PSD was estimated for C3 and C4 in each EEG epoch using Welch’s method. Frequencies from 0.5 to 30 Hz were retained, concatenated across channels, and log-transformed. Inputs and targets were standardized dimension by dimension using means and standard deviations from valid training-set EEG epochs. A Transformer matching the temporal architecture of the main model forecast the next 10 min of PSD from the preceding 30 min. The model was trained using mean absolute error and selected on validation performance. Forecast PSD was decoded by an MLP applied to each EEG epoch and fitted exclusively to true PSD in the validation set.

### Input-history length and forecast performance

To compare the effects of input-history length and forecast horizon without training a model for each combination, we used one set of weights across all length combinations. A separate variable-length Transformer sequence predictor with rotary positional encoding (RoPE) [42] was trained using the same frozen EEG epoch encoder and data split. Predictor weights were initialized from the primary fixed-window checkpoint. Input-history length H and forecast horizon F both used the grid {0.5, 1, 2, 3, 4, 5, 10, 15, …, 360} min, with H + F ≤ 480 min, yielding 4,753 combinations.

During testing, all eligible windows were enumerated independently at 30-s intervals for each combination.

### Target-directed integrated gradients

Integrated gradients (IG) [13] were used to quantify the contributions of historical EEG-frame CNN features to forecast proximity to a target sleep stage. Attribution analysis used the frozen EEG epoch encoder and the primary sequence predictor.

We fitted principal component analysis (PCA) to the two-dimensional latent states in the validation set. We held the center, loadings, component signs and standard deviations fixed when transforming observed and forecast latent states in the test set using *T* (*z*)=[(*z* − *μ*_*val*_)*V* _*val*_]*⊘ σ* _*val*_. The target centroid *c*_*s*_ was the mean of all valid latent states belonging to the corresponding stage in the test set. The NREM centroid pooled N1/N2/N3 epochs. This target geometry was used for interpretive analysis of the frozen model.

The objective was the negative mean distance from the 10 forecast latent states in the first 5 min to the target centroid:

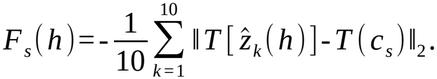

We defined the forecast start as the first epoch after a manually scored stage transition, with 30 min of input and 10 min of forecasting within the same night. The source and target stages each had to persist for at least 5 min before and after the transition, respectively. All epochs within the complete window had to be valid. The analysis covered eight directions: N2⟶N3/REM/Wake, N3⟶N2/REM/Wake, and REM⟶NREM/Wake. REM⟶NREM pooled REM⟶N1/N2/N3 events that individually met these criteria. Candidate selection was independent of forecast outcomes.

Integrated gradients were evaluated at 64 points along the straight path from an all-zero CNN feature baseline to the observed historical features. Model parameters remained frozen, and actual future EEG was not supplied to the predictor. Summing across feature dimensions yielded frame-level attribution 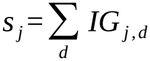. Positive and negative values indicated positive and negative contributions to target proximity relative to the zero baseline. Attribution completeness was assessed by comparing the sum of all attributions with the objective change from baseline to observed input. No completeness threshold was used to exclude windows. We also selected up to 4 windows per direction at evenly spaced positions in candidate order, yielding 32 windows. Calculations were repeated with 256 points to compare frame-attribution rankings, sets of high-contribution frames, and sign agreement.

Subsequent spectral and transition-direction axis analyses used high-contribution frames from the final 5 min of input. These were defined as frames with absolute attribution at or above the window-specific 75th percentile and grouped by attribution sign. Windows containing at least 3 frames in each sign group were included.

### Attribution-related EEG spectra

Relative power spectral density (PSD) was calculated separately for high-contribution frames with positive and negative attributions in the final 5 min of input. For each 3-s frame, C3/C4 power spectra were estimated using Welch’s method with a Hann window. The two channels were averaged, and the resulting spectrum was normalized by its integral over 0.5–30 Hz. Within each window, relative PSD was averaged separately across positive and negative frames, and the difference between their log_10_ values was calculated.

To test whether relative PSD differed systematically between positive- and negative-IG frames within each transition direction, window-level log10 PSD differences were first averaged within each night. Frequency-wise inference used a two-sided, night-level sign-flipping maximum cluster-mass test with 10,000 permutations and an uncorrected cluster-forming threshold of *P* < 0.05 [43]. Permutation maxima controlled the family-wise error rate within each direction. Frequency bands were delta (0.5–4 Hz), theta (4–8 Hz), alpha (8–12 Hz), sigma (12–15 Hz), and beta (15–30 Hz).

### Transition-direction axes and counterfactual perturbations

A transition-direction axis was defined by the difference between the mean standardized CNN features of positive- and negative-IG high-contribution frames for that direction, normalized to unit length. For each event, the difference between mean standardized features of positive and negative frames was calculated as 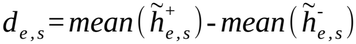 Event-level differences were then averaged with equal weight and normalized to unit length, yielding 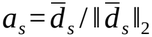

To assess reproducibility across nights and target specificity, five-fold cross-validation was performed with folds defined by night. For each fold, feature means, standard deviations and axes were estimated only from the other four folds; feature standardization weighted nights equally. High-contribution frames from held-out nights were projected onto the estimated transition-direction axes. The AUC for distinguishing positive from negative IG frames was calculated per night and summarized with equal weight across nights. Matched-axis AUC was compared with 0.5. Using the same held-out frames and sign labels, we also compared it with the mean AUC of other target axes sharing the same source stage. The two families of comparisons were each corrected across the eight directions using the Holm procedure.

To test whether changing features along transition-direction axes shifted forecasts toward the target stage, we applied within-model perturbations in stable sleep windows. Stable windows required the same stage throughout the 5 min before and 5 min after the forecast start. Experiments across perturbation magnitudes used common axes and standard deviations re-estimated from all eligible test transitions. Nights receiving perturbations could also contribute to axis estimation. We applied 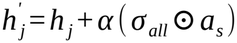 to all CNN frame features in the final 5 min of input, with α = 0, 1, …, 15. All other historical features were unchanged, and trajectories were generated through the remaining frozen encoder layers and sequence predictor. Here, α is the displacement length along a unit axis in standardized feature space.

To test whether IG-related spectral changes induced corresponding forecast shifts at the raw-signal level, we constructed spectral perturbation templates for each direction. For each direction, window-level differences in log_10_ relative PSD between positive and negative attribution frames were averaged within nights and then equally across test nights. The resulting profiles were smoothed to obtain spectral templates. All recipient nights used the same template, and recipient nights were not excluded from template estimation. In the same stable windows, we Fourier-transformed C3/C4 for each 30-s EEG epoch in the final 5 min of input. Phase was preserved, and amplitudes were adjusted according to 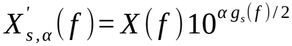, with α = 0, 0.5, …, 5. After inverse transformation, the perturbed signals were passed through the frozen model. Actual PSD, waveform correlations, and root mean square (RMS) ratios were also recorded.

The α values for CNN and PSD perturbations refer to different perturbation spaces.

