## Supplemental Data 1 for "Forecasting Human Sleep Dynamics with a Latent-State Framework"

### Supplementary Materials

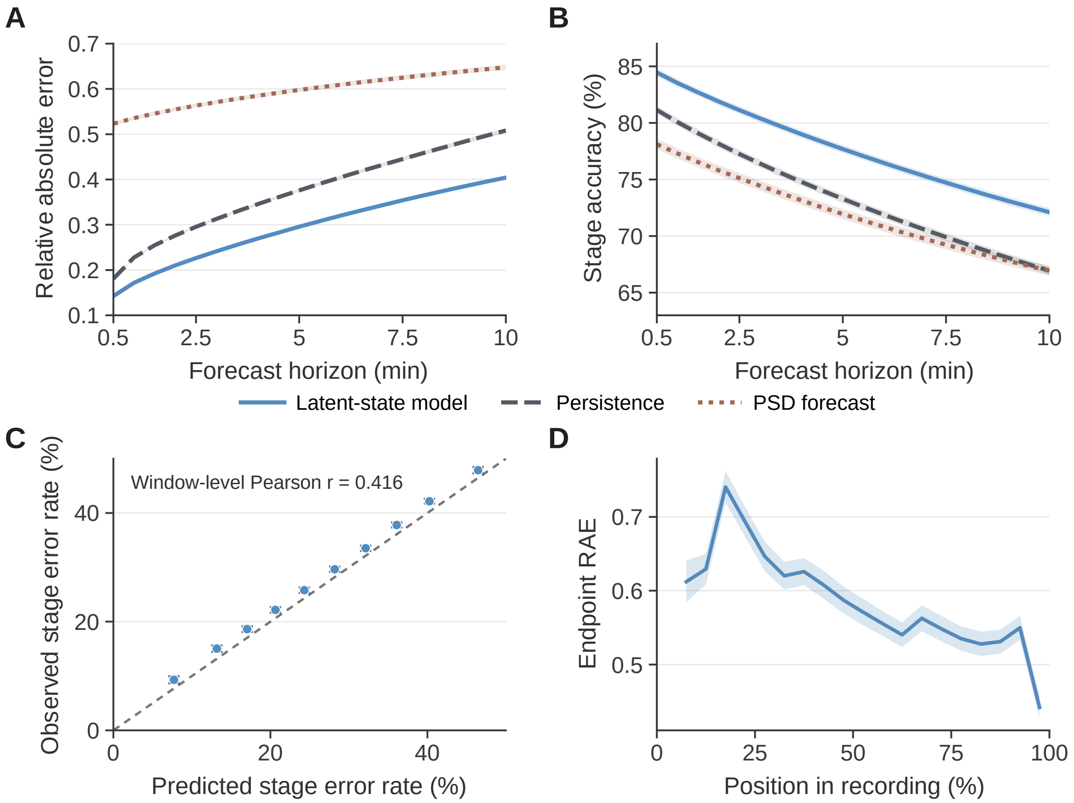


**Fig. S1. Sleep forecasting performance, predicted error and variation across the night.** (A) Relative absolute error (RAE) and (B) four-class sleep-stage accuracy over the indicated forecast horizon in 1,084 held-out SHHS nights. The model used 30 min of EEG history to forecast the next 10 min. The persistence baseline carried the last observed state forward. RAE was calculated on the latent-state scale for the model and persistence, and on the log power spectral density (PSD) scale for the spectral baseline. At 10 min, mean RAE was 0.404, 0.508 and 0.648, and accuracy was 72.11%, 66.94% and 66.93% for the latent-state model, persistence and spectral forecasting, respectively. (C) Predicted and observed four-class sleep-stage error rates over 10 min. Windows were ranked by the mean predicted error probability across 20 epochs and divided into deciles. Observed error was the fraction of epochs misclassified against manual staging. The dashed line marks equality. The lowest-risk decile had an observed error rate of 9.29%. Across all 916,633 windows, Pearson r = 0.416. (D) RAE at the 10-min endpoint by relative position in the full recording, including Wake. Each bin spans 5% of the recording. The first bin contained no eligible endpoints. Mean endpoint RAE was 0.6428 in the first half and 0.5363 in the second (paired difference, −0.1065, 95% CI −0.1148 to −0.0983). The confidence interval for the difference was obtained from 10,000 bootstrap resamples of nights, keeping the two halves of each night paired. Plotted means average windows within each contributing night and weight nights equally. Shading and vertical error bars show pointwise 95% confidence intervals from night-level bootstrap resampling

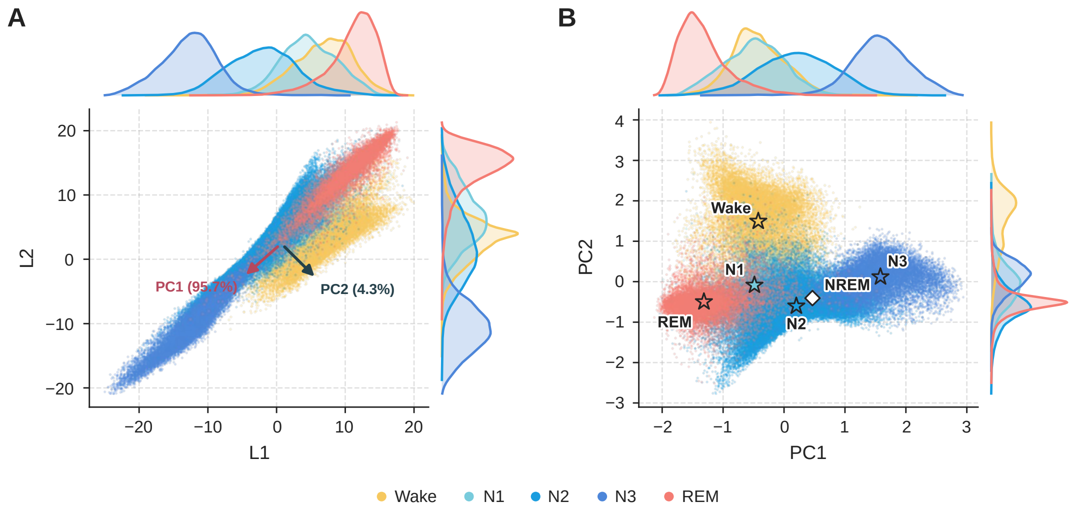


**Fig. S2. Sleep-stage distributions and centroids in the EEG representation space.** (A) Each point represents one of 100,000 valid 30-s epochs sampled uniformly without replacement from 1,084 held-out SHHS nights. Color indicates the scored sleep stage. L1 and L2 are the two encoder coordinates. Arrows show the principal-component directions fitted to 600,713 valid epochs from 650 validation nights. PC1 and PC2 accounted for 95.7% and 4.3% of validation-set variance, respectively. (B) The same epochs after projection onto the two principal components and division by their validation-set standard deviations. Stars mark the centroids of Wake, N1, N2, N3 and REM. Each centroid is the mean over all valid test epochs scored in that stage. The open diamond marks the pooled NREM centroid used for the REM→NREM direction. Marginal curves in both panels show the within-stage densities along each axis.

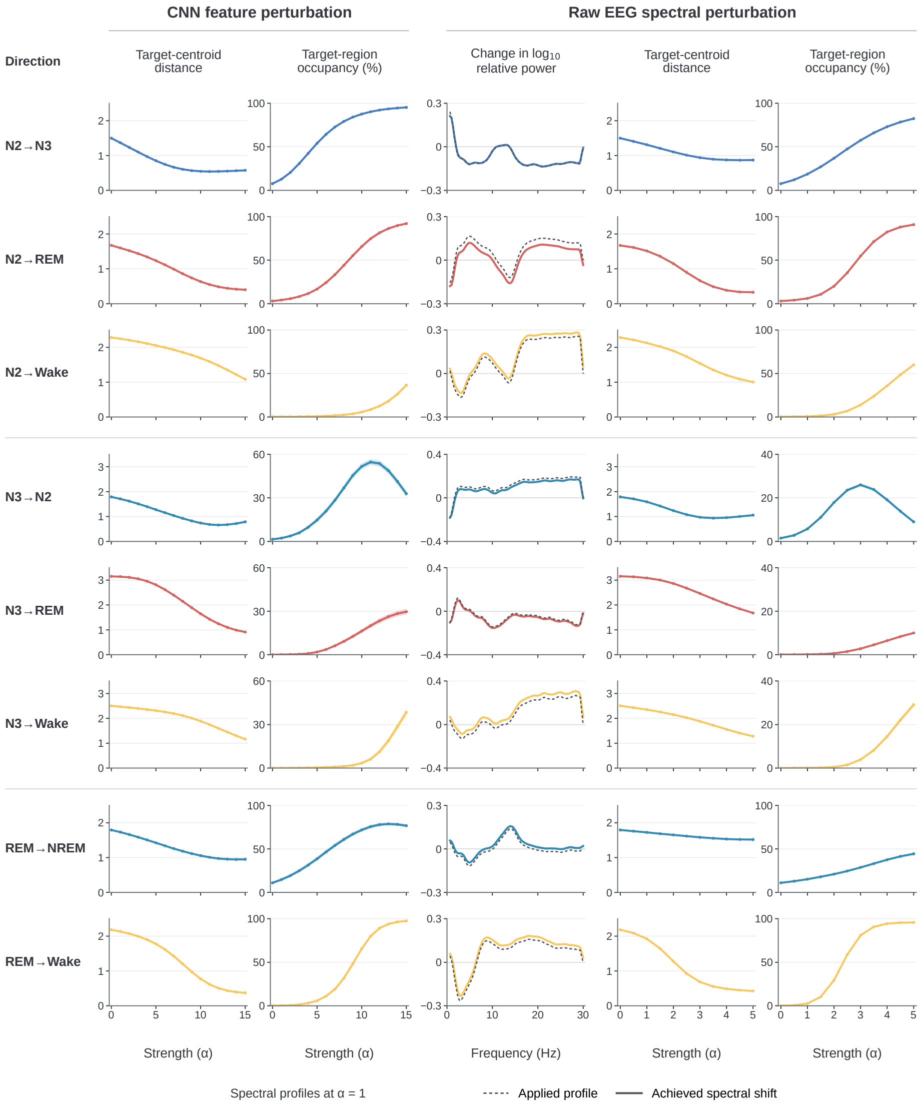


**Fig. S3. Responses to CNN feature and EEG spectral perturbations in stable sleep.** Rows show eight perturbation directions. The scored stage was stable for 5 min on either side of the forecast origin (N2, 116,706 windows from 1,065 nights; N3, 38,413 from 737; REM, 51,944 from 1,011). Perturbations were applied to the final 5 min of input along a CNN feature axis (α = 0–15) or by reshaping the raw EEG spectrum while preserving Fourier phase (α = 0–5). The middle column compares the applied and achieved changes in $\log_{10}$ relative power at α = 1. The first 5 min of the forecast were summarized by the mean distance to the target centroid in standardized PC1–PC2 space and the percentage of epochs nearest the target centroid. For REM→NREM, distance uses the pooled NREM centroid and occupancy includes N1, N2 and N3. Nights were weighted equally. Shading shows 95% night-bootstrap intervals for CNN responses (2,000 resamples) and ±1 s.e.m. for achieved spectra and raw-EEG responses. For N3→N2, occupancy peaked at 54.7% (α = 11) and 26.0% (α = 3), and distance reached minima of 0.657 (α = 12) and 0.935 (α = 3.5), for CNN and spectral perturbations, respectively.

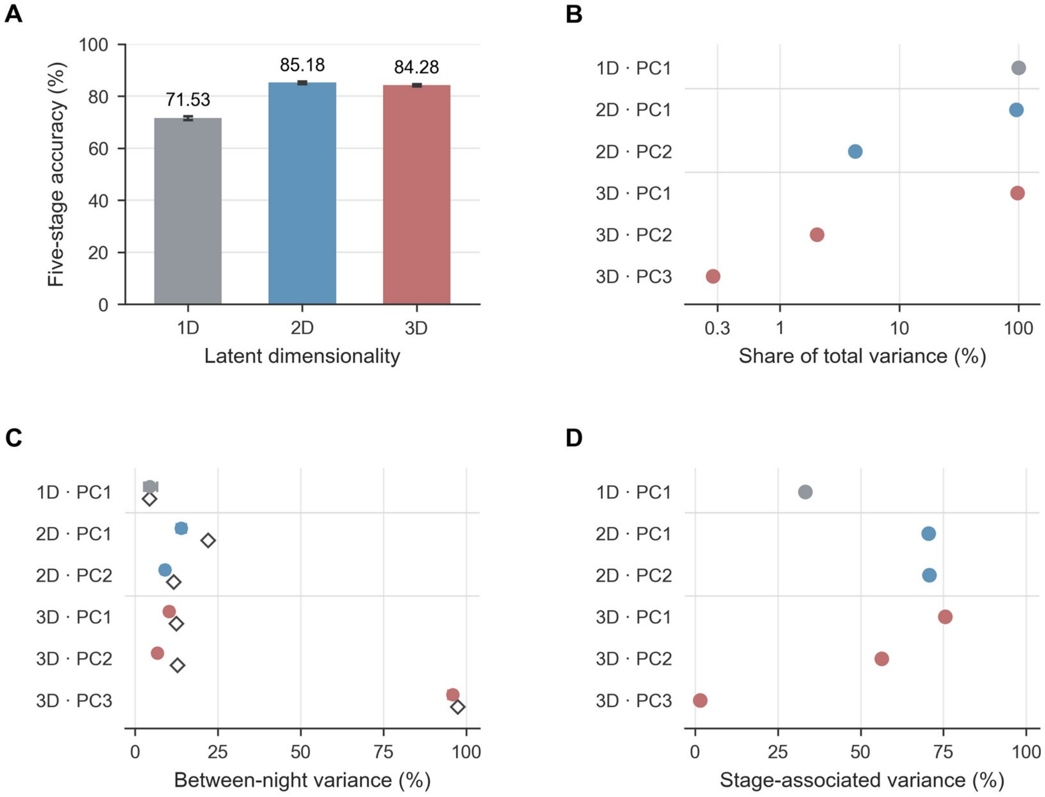


**Fig. S4. Sleep-stage information and variance structure across latent dimensionalities.**(A) Current-epoch five-stage accuracy for matched encoders with one, two or three latent dimensions. Each bar is an equally weighted mean across the same 650 validation nights. (B-D) Principle components (PCs) were fitted separately for each encoder to 600,713 valid 30-s epochs. Each epoch carries equal weight in these variance estimates. (B) Fraction of total latent variance explained by each PC, on a logarithmic scale. (C) Fraction of each PC's variance attributable to differences between nightly means (filled circles). Open diamonds give the fraction after subtracting stage-specific mean representations while holding the PC directions fixed. (D) Fraction of each PC's variance explained by differences between the means of the five manually scored stages ($\eta^2$). In the two-dimensional encoder, PC2 carried 4.28% of total variance and 70.72% of its variance was stage-associated. In the three-dimensional encoder, PC3 carried 0.274% of total variance and 95.92% of its variance was between nights. Error bars in A and for filled circles in C indicate pointwise 95% percentile confidence intervals from 10,000 night-level bootstrap resamples, conditional on the fitted models and PC directions. One encoder was evaluated per dimensionality.

**Table S1. Discrimination of sustained sleep-stage transitions from continued stable sleep.**

| Transition | Nights | Mean night-level AUC | 95% CI |
|---|---|---|---|
| Overall | 995 | 0.895 | 0.884–0.906 |
| N2 → N3 | 327 | 0.980 | 0.972–0.987 |
| N2 → REM | 703 | 0.975 | 0.967–0.983 |
| N2 → Wake | 431 | 0.758 | 0.729–0.787 |
| N3 → N2 | 164 | 0.905 | 0.868–0.939 |
| N3 → REM | 37 | 0.987 | 0.958–1.000 |
| N3 → Wake | 70 | 0.765 | 0.688–0.837 |
| REM → NREM | 408 | 0.956 | 0.943–0.968 |
| REM → Wake | 197 | 0.678 | 0.627–0.727 |

The model used 30 min of EEG to forecast the next 10 min. Each transition forecast began at the first epoch of the target stage, with at least 5 min in each adjacent stage. Stable windows remained in the same source stage for the final 5 min of input and all 10 min of the forecast. The score was the mean predicted probability of the target over the 20 forecast epochs. The N2 target used the N1/N2 probability, and NREM used the sum of the N1/N2 and N3 probabilities. AUC is the area under the receiver operating characteristic curve. Each direction includes nights with at least one transition and one same-source stable window. Nightly AUCs were averaged with equal weight. For the overall estimate, available directions were averaged within each night before averaging across nights. The 95% confidence intervals (CIs) are percentile intervals from 10,000 whole-night bootstrap resamples.

**Table S2. Discrimination of positive- and negative-IG frames by individual CNN feature axes.**

Rows identify the transition from which frames were selected by integrated gradients (IG). Columns identify the convolutional neural network (CNN) feature axis used to score those frames. Bold entries mark the matched axis. Values are the mean area under the receiver operating characteristic curve (AUC) ± standard error across the indicated nights. In fivefold cross-validation grouped by night, AUC was calculated separately for each held-out night with positive-IG frames as the positive class. Each night contributed equally. An AUC of 0.5 indicates chance discrimination. Two-sided Wilcoxon signed-rank tests compared nightly matched-axis AUCs with 0.5 and, separately, with the same nights' mean AUC for alternative axes from the same source stage. Holm correction was applied across the eight directions separately for each comparison. The matched axis yielded the highest mean AUC for each of the eight transitions. The N2→REM axis scored N2→N3 frames with a mean AUC of 0.209, and the N2→N3 axis scored N2→REM frames with a mean AUC of 0.182. In both cases, positive-IG frames tended to receive lower scores than negative-IG frames.

**A. N2 source stage**

| IG-frame transition | Nights | N2→N3 | N2→REM | N2→Wake |
|---|---|---|---|---|
| N2→N3 | 300 | **0.842 ± 0.007** | 0.209 ± 0.008 | 0.371 ± 0.009 |
| N2→REM | 731 | 0.182 ± 0.004 | **0.852 ± 0.004** | 0.688 ± 0.005 |
| N2→Wake | 438 | 0.412 ± 0.008 | 0.621 ± 0.009 | **0.840 ± 0.005** |

**B. N3 source stage**

| IG-frame transition | Nights | N3→N2 | N3→REM | N3→Wake |
|---|---|---|---|---|
| N3→N2 | 202 | **0.813 ± 0.008** | 0.671 ± 0.010 | 0.485 ± 0.011 |
| N3→REM | 41 | 0.713 ± 0.019 | **0.818 ± 0.017** | 0.672 ± 0.019 |
| N3→Wake | 60 | 0.528 ± 0.024 | 0.588 ± 0.021 | **0.794 ± 0.015** |

**C. REM source stage**

| IG-frame transition | Nights | REM→NREM | REM→Wake |
|---|---|---|---|
| REM→NREM | 446 | **0.782 ± 0.007** | 0.605 ± 0.007 |
| REM→Wake | 233 | 0.707 ± 0.011 | **0.840 ± 0.009** |

**Table S3. Spectral differences between positive- and negative-IG EEG frames. A. Band-level effects.**

Relative power was integrated over each band in the mean spectrum of the selected frames of each IG sign. Within-window differences in $\log_{10}$ relative band power (positive minus negative IG) were averaged within each night and then across nights with equal weight. The 95% confidence intervals are unadjusted percentile intervals from 10,000 night-level bootstrap resamples. Two-sided Wilcoxon signed-rank tests against zero were Holm-adjusted across the 40 direction-by-band comparisons.

| Transition | Band (Hz) | Nights | Mean effect | 95% CI | Holm-adjusted P |
|---|---|---|---|---|---|
| N2→N3 | Delta (0.5–4) | 300 | 0.0622 | 0.0563 to 0.0682 | $7.53 \times 10^{-41}$ |
| N2→N3 | Theta (4–8) | 300 | −0.1337 | −0.1454 to −0.1216 | $5.77 \times 10^{-42}$ |
| N2→N3 | Alpha (8–12) | 300 | −0.0633 | −0.0789 to −0.0476 | $6.51 \times 10^{-12}$ |
| N2→N3 | Sigma (12–15) | 300 | −0.0043 | −0.0236 to 0.0153 | 1.000 |
| N2→N3 | Beta (15–30) | 300 | −0.1348 | −0.1540 to −0.1156 | $1.35 \times 10^{-28}$ |
| N2→REM | Delta (0.5–4) | 731 | −0.0385 | −0.0439 to −0.0332 | $1.22 \times 10^{-39}$ |
| N2→REM | Theta (4–8) | 731 | 0.1361 | 0.1301 to 0.1422 | $2.37 \times 10^{-116}$ |
| N2→REM | Alpha (8–12) | 731 | 0.0399 | 0.0312 to 0.0483 | $9.10 \times 10^{-17}$ |
| N2→REM | Sigma (12–15) | 731 | −0.1152 | −0.1281 to −0.1019 | $9.37 \times 10^{-49}$ |
| N2→REM | Beta (15–30) | 731 | 0.0828 | 0.0711 to 0.0945 | $1.26 \times 10^{-36}$ |
| N2→Wake | Delta (0.5–4) | 438 | −0.0747 | −0.0835 to −0.0657 | $5.64 \times 10^{-39}$ |
| N2→Wake | Theta (4–8) | 438 | 0.0103 | −0.0015 to 0.0218 | 0.00387 |
| N2→Wake | Alpha (8–12) | 438 | 0.0896 | 0.0759 to 0.1030 | $7.82 \times 10^{-31}$ |
| N2→Wake | Sigma (12–15) | 438 | −0.0473 | −0.0657 to −0.0292 | $5.07 \times 10^{-4}$ |
| N2→Wake | Beta (15–30) | 438 | 0.2233 | 0.2047 to 0.2419 | $6.77 \times 10^{-60}$ |
| N3→N2 | Delta (0.5–4) | 202 | −0.0463 | −0.0561 to −0.0380 | $1.05 \times 10^{-23}$ |
| N3→N2 | Theta (4–8) | 202 | 0.1017 | 0.0832 to 0.1194 | $3.97 \times 10^{-17}$ |
| N3→N2 | Alpha (8–12) | 202 | 0.0897 | 0.0659 to 0.1133 | $3.81 \times 10^{-11}$ |
| N3→N2 | Sigma (12–15) | 202 | 0.1132 | 0.0914 to 0.1346 | $6.37 \times 10^{-16}$ |
| N3→N2 | Beta (15–30) | 202 | 0.1866 | 0.1542 to 0.2189 | $1.45 \times 10^{-19}$ |

**Table S3. Spectral differences between positive- and negative-IG EEG frames (continued).**

**A. Band-level effects.**

Effect sizes are night-level means in $\log_{10}$ relative-power units. Positive values indicate higher power in positive-IG frames. The 40 comparisons share one Holm correction.

| Transition | Band (Hz) | Nights | Mean effect | 95% CI | Holm-adjusted P |
|---|---|---|---|---|---|
| N3→REM | Delta (0.5–4) | 41 | 0.0095 | −0.0028 to 0.0212 | 0.504 |
| N3→REM | Theta (4–8) | 41 | −0.0053 | −0.0475 to 0.0371 | 1.000 |
| N3→REM | Alpha (8–12) | 41 | −0.1209 | −0.1760 to −0.0649 | 0.0025 |
| N3→REM | Sigma (12–15) | 41 | −0.0658 | −0.1140 to −0.0146 | 0.0648 |
| N3→REM | Beta (15–30) | 41 | −0.0433 | −0.0998 to 0.0164 | 0.504 |
| N3→Wake | Delta (0.5–4) | 60 | −0.0139 | −0.0311 to 0.0025 | 1.000 |
| N3→Wake | Theta (4–8) | 60 | −0.0464 | −0.0793 to −0.0124 | 0.225 |
| N3→Wake | Alpha (8–12) | 60 | 0.0118 | −0.0536 to 0.0809 | 1.000 |
| N3→Wake | Sigma (12–15) | 60 | 0.0237 | −0.0476 to 0.0951 | 1.000 |
| N3→Wake | Beta (15–30) | 60 | 0.2375 | 0.1351 to 0.3436 | 0.0168 |
| REM→NREM | Delta (0.5–4) | 446 | −0.0070 | −0.0143 to 0.0003 | 1.000 |
| REM→NREM | Theta (4–8) | 446 | −0.0815 | −0.0886 to −0.0747 | $1.08 \times 10^{-57}$ |
| REM→NREM | Alpha (8–12) | 446 | 0.0075 | −0.0019 to 0.0167 | 0.738 |
| REM→NREM | Sigma (12–15) | 446 | 0.1311 | 0.1165 to 0.1453 | $2.26 \times 10^{-46}$ |
| REM→NREM | Beta (15–30) | 446 | 0.0267 | 0.0128 to 0.0407 | 0.00568 |
| REM→Wake | Delta (0.5–4) | 233 | −0.0940 | −0.1083 to −0.0802 | $1.92 \times 10^{-24}$ |
| REM→Wake | Theta (4–8) | 233 | −0.0610 | −0.0755 to −0.0467 | $1.84 \times 10^{-13}$ |
| REM→Wake | Alpha (8–12) | 233 | 0.1466 | 0.1276 to 0.1659 | $4.26 \times 10^{-29}$ |
| REM→Wake | Sigma (12–15) | 233 | 0.0986 | 0.0806 to 0.1164 | $1.50 \times 10^{-18}$ |
| REM→Wake | Beta (15–30) | 233 | 0.1398 | 0.1209 to 0.1592 | $4.37 \times 10^{-27}$ |

**Table S3. Spectral differences between positive- and negative-IG EEG frames (continued).**
**B. Significant frequency clusters.**
Positive and negative IG indicate which frames had higher relative power within the cluster. Frequencies are the first and last sampled bins in each significant cluster. Two-sided night-level sign flips (10,000 permutations) were used with a cluster-forming threshold of $P < 0.05$. Cluster mass is the sum of absolute $t$ statistics across adjacent bins with the same sign. The maximum cluster mass across frequencies controlled the family-wise error rate within each transition direction.

| Transition | Frequency (Hz) | Higher relative PSD | Nights | Cluster mass | FWER-adjusted P |
|---|---|---|---|---|---|
| N2→N3 | 0.67–1.67 | Positive IG | 300 | 80.80 | 0.0012 |
| N2→N3 | 2.00–10.00 | Negative IG | 300 | 253.99 | $1.00 \times 10^{-4}$ |
| N2→N3 | 14.33–30.00 | Negative IG | 300 | 436.64 | $1.00 \times 10^{-4}$ |
| N2→REM | 0.67–1.67 | Negative IG | 731 | 95.75 | 0.0011 |
| N2→REM | 2.00–10.67 | Positive IG | 731 | 586.84 | $1.00 \times 10^{-4}$ |
| N2→REM | 11.33–15.00 | Negative IG | 731 | 123.35 | $3.00 \times 10^{-4}$ |
| N2→REM | 15.67–30.00 | Positive IG | 731 | 697.10 | $1.00 \times 10^{-4}$ |
| N2→Wake | 1.00–5.00 | Negative IG | 438 | 186.75 | $1.00 \times 10^{-4}$ |
| N2→Wake | 6.67–11.00 | Positive IG | 438 | 129.59 | 0.0013 |
| N2→Wake | 12.67–14.33 | Negative IG | 438 | 30.76 | 0.0224 |
| N2→Wake | 15.00–30.00 | Positive IG | 438 | 867.80 | $1.00 \times 10^{-4}$ |
| N3→N2 | 0.67–1.33 | Negative IG | 202 | 49.28 | 0.0089 |
| N3→N2 | 1.67–30.00 | Positive IG | 202 | 682.61 | $1.00 \times 10^{-4}$ |
| N3→REM | 0.67–1.33 | Negative IG | 41 | 16.62 | 0.0439 |
| N3→REM | 1.67–3.00 | Positive IG | 41 | 32.56 | 0.0182 |
| N3→REM | 8.33–12.67 | Negative IG | 41 | 54.87 | 0.0081 |
| N3→REM | 27.67–30.00 | Negative IG | 41 | 22.46 | 0.03 |
| N3→Wake | 1.33–6.33 | Negative IG | 60 | 61.87 | 0.0108 |
| N3→Wake | 15.00–30.00 | Positive IG | 60 | 174.85 | $2.00 \times 10^{-4}$ |
| REM→NREM | 0.67–1.33 | Positive IG | 446 | 15.88 | 0.0416 |
| REM→NREM | 1.67–9.33 | Negative IG | 446 | 232.70 | $1.00 \times 10^{-4}$ |
| REM→NREM | 10.67–17.00 | Positive IG | 446 | 187.81 | $1.00 \times 10^{-4}$ |
| REM→Wake | 1.67–6.33 | Negative IG | 233 | 250.81 | $2.00 \times 10^{-4}$ |
| REM→Wake | 7.00–30.00 | Positive IG | 233 | 659.65 | $1.00 \times 10^{-4}$ |
